# A Quest for a Histaminergic or Orexinergic Biomarker for Sudden Infant Death Syndrome

**DOI:** 10.1101/2025.10.08.680830

**Authors:** Yan Zhao, Guang-Fu Cui, Sabine Plancoulaine, Marion Comajuan, Marine Bersault, Clara Inocente, Yi-Ping Hou, Yves Dauvilliers, Lotierzo Manuela, Aurore Guyon, David Meyronet, Anne Jouvet, Beatrice Kugener, Marine Thieux, Philippe Robert, Jian-Sheng Lin, Patricia Franco

## Abstract

**Aims:** This study aimed to investigate the role of the Hypocretin/Orexin (Ox) and Histamine (HA) systems—two key regulators of wakefulness —in sudden infant death syndrome (SIDS), a condition characterized by impaired arousal responses during sleep.

**Methods:** Cerebrospinal fluid (CSF) Ox levels were measured in 61 healthy controls, 70 Sudden Unexpected Death Infants (SUDI) (38 SIDS cases and 32 explained deaths (ED). HA and its metabolite, tele-methylhistamine (t-MeHA), were analysed in an additional 46 SUDI (34 SIDS, 12 ED) and 42 controls. Immunocytochemistry was performed on hypothalamic tissue from 11 SIDS and 13 ED cases to assess the number of Ox and HA neurons, using markers for Orexin-A and histidine decarboxylase (HDC), respectively.

**Results:** Overall CSF Ox levels did not differ significantly between groups. However, while an age-related decline in Ox levels was observed in controls (0–2 vs. 2–6 months), this pattern was absent in deceased infants, resulting in relatively higher Ox concentrations in the 2–6 month age group. HA and t-MeHA levels were significantly elevated in both SIDS and ED cases, likely due to postmortem release.

Immunohistochemical analysis revealed an increased number of Ox-immunoreactive neurons in the rostral and caudal dorsolateral hypothalamus in SIDS cases compared to EDs. No differences were observed in the number or morphology of HA neurons.

**Conclusion:** Findings suggest increased Ox activity in SIDS cases, potentially reflecting repeated exposure to stress or hypoxia prior to death. In contrast, HA neurons do not appear to play a role in the pathophysiology of SIDS. Further research is needed to identify more specific biomarkers for assessing SIDS risk.

## 1. INTRODUCTION

Sudden infant death syndrome (SIDS) is defined as the death of an infant under one year of age which occurs during the sleep and remains unexplained after a complete evaluation of death scene, clinical history and postmortem investigation. To date, SIDS remains the leading cause of post-neonatal mortality (between one month and one year)[1, 2]. Epidemiological studies have enabled a better understanding of SIDS risk factors, including, prone sleep position, bed-sharing, overheating, infections, prenatal or postnatal cigarette exposure, prematurity and low birth weight[1]. Thus, SIDS incidence has dramatically decreased over recent decades thanks to awareness campaigns[2, 3].

Nevertheless, the physiopathology of SIDS is still poorly understood. Its occurrence during sleep at a specific developmental window, especially between 2 and 6 months of age, suggests an association with a dysfunctional maturation of sleep-wake regulation especially within the brain’s arousal systems. Polysomnographic recordings have shown that SIDS victims exhibit altered arousability, including fewer arousals at the end of the night and an increased prevalence of incomplete arousals compared to age matched healthy infants [4–6]. However, the specific arousal system(s) involved remains unknown.

The orexinergic and histaminergic systems, both located in the posterior hypothalamus play a distinctive yet synergetic role in promoting wakefulness. Dysfunction in either system may impair arousal. Orexin (Ox, also called as hypocretin) has been involved in sleep disorders such as narcolepsy[7]. The Ox system undergoes a maturation process during early infancy, characterized by a progressive increase in cerebrospinal fluid (CSF) Ox levels during the late prenatal period and the first months after birth, followed by a decline around the age of peak SIDS incidence (2–6 months) [8, 9]. In our previous evaluation of CSF Ox in healthy children between 0 to 18 years, we found that infants aged 2 to 6 months had lower CSF Ox levels than those in other age groups[8]. This period of low Ox levels coincides with the peak incidence of SIDS. Moreover, decreased Ox immunoreactivity has been found in both the hypothalamus and brainstem of SIDS victims [10], supporting the potential involvement of the Ox system in the physiopathology of SIDS.

These findings also raise the question of whether histamine (HA) neurons may be implicated during the critical period of SIDS, given the close anatomical and functional interactions between the orexin (OX) and HA systems[11, 12]. Impaired HA transmission has been associated with reduced wakefulness and increased sleepiness in both animal models and human cases of childhood narcolepsy[12–14]. However, findings in adults with hypersomnia have been inconsistent [15]. In a recent large-scale study of healthy individuals from birth to adulthood[16], we found elevated cerebrospinal fluid (CSF) levels of HA and its direct metabolite, tele-methylhistamine (t-MeHA), during the first six months of life compared to older age groups. This suggests that enhanced HA transmission in early infancy may play a protective role against SIDS.

Therefore, this study aims to investigate whether the Ox and HA systems are altered in SIDS victims by addressing the following objectives: 1) To determine whether CSF Ox and HA levels differ between healthy infants and SIDS victims and infants who died of other causes (ED) across three development age groups (0-2, 2-6 and 6-12 months); 2) to assess whether the number of Ox or HA neurons is altered in SIDS cases by immunohistochemical staining using antibodies against Ox and histidine decarboxylase, the enzyme responsible for HA synthesis.

## 2. PATIENTS AND METHODS

### 2.1. First part: CSF study

#### 2.1.1. CSF Orexin levels Patients

CSF samples for Ox measurements were collected from 61 control infants and 70 Sudden Unexpected Death Infants (SUDI) (38 SIDS cases and 32 infants died from known causes (ED)) at the Hôpital Femme Mère Enfant in Lyon, France.

For SUDI children, clinical information included: circumstances of death, delay between CSF puncture and death, known risk factors of SIDS (ventral position (n=10), co-sleeping (n=8), smoke exposure (n=25), infection (n=27). SIDS were included only if no explanation to the death was found after the regular clinical and paraclinical examinations and autopsy. Five SIDS infants (13.2%) were born prematurely (median 32.5 Gestational Age (GA) weeks (range from 29 to 36 weeks). Thirty-two infants died from known causes (cardiopathy (n=5), fibroelastosis (n=1), pericarditis (n=1), asphyxia (n=12), sepsis (n=1), meningitis (n=1), pulmonary infectious (n=1), epileptic encephalopathy (n=1), child abuse (n=4), inhalation (n=1), intoxication (n=1), traumatism (n=1), volvulus (n=1), gastro-enteritis (n=1). Nine (29%) of these infants were born prematurely (median 34 weeks GA (range from 27 to 36 weeks)).

Control CSF were collected from infants below 1 year old for meningitis suspicion. During a lumbar puncture, a spare tube was usually collected for possible further analyses. We used this extra CSF sample as a control if it was not clinically useful anymore and if the results of CSF analysis were normal (cell count, glucose, proteins and lactic acid). For each infant, clinical information was collected: date and time of lumbar puncture, date of birth, delivery and birth circumstances, medical background and reason of the lumbar puncture. No control infants had a preterm delivery.

Most lumbar punctures were performed to rule out meningitis in a context of fever; final diagnosis included: acute pyelonephritis (n=13), gastroenteritis (n=3), bronchiolitis (n=5), viral meningitis (n=4), other infections (n=4) and unexplained fever/virosis (n= 21). Other lumbar punctures were performed for epilepsy (n=1), Hirschsprung disease (n=1), portal cavernoma (n=1), bad weight growth (n=3), bulging fontanel (n=1), without access to the medical file for 4 children.

##### Orexin measurements

CSF were stored at -80°C. Hypocretin-1 (orexin-A) was measured from CSF samples as previously described in Gui-de-Chauliac Hospital in Montpellier, France[17]. It was also determined in all the infants in duplicate from CSF samples without prior extraction using ^125^I radioimmunoassay kits from Phoenix Peptide, Inc, according to manufacturer guidelines. The detection limit was 10 pg/ml and intra-assay variability was less than 10%. All values were back-referenced to Stanford reference samples (HHMI Stanford University Center for Narcolepsy, Palo Alto CA).

#### 2.1.2. CSF Histamine levels Patients

The CSF samples for HA values were collected from 46 other SUDI infants (34 SIDS and 12 ED) and 42 other controls at the Hôpital Femme Mère Enfant in Lyon, France. There were no preterm babies in this group of control infants. Indications for lumbar puncture were available for N=30 and were fever (N=24), febrile seizures (N=2), acute gastroenteritis with dehydration (N=2), respiratory distress by meconium. All biological results were subsequently found to be normal.

##### Measurements of histamine and its direct metabolite

The HA and t-MeHA assays were carried out at Bioproject-Biotech in Saint Gregoire (France). The technique used is the one developed by Croyal *et al.* [18] for the simultaneous analysis of CSF HA and its main stable metabolite t-MeHA. All measurements were performed blind. The CSF samples were immediately preserved at -80° C after sampling until use. Blood contamination was checked visually and specimens of the CSF with abnormal color were excluded from the study. These measurements rely on derivatization of primary amines using 4-bromobenzenesulfonyl chloride and subsequent analysis by reverse liquid chromatography (UPLC: ultra-performance liquid chromatography) with mass spectrometric detection[18].

### 2.2. Second part: Study on orexin and histamine neurons by immunohistochemistry

#### 2.2.1. Postmortem brain material

Formalin-fixed, paraffin-embedded posterior hypothalamus (11 SIDS and 8 ED) were collected from the pathological department of the Hôpital Femme Mère Enfant in Lyon, France. In SIDS, there were 2 preterm babies (27 and 30 weeks GA), 6/11 (54.5%) were boys and the median age was 3.5 months (range 1-9 months). In deceased controls, there were 4 preterm babies (33, 34, 34 and 36 weeks), 4/8 were boys (50%) and the median age was 4.5 months (range from 0.5-7 months).

##### Ox and HDC immunohistochemistry

The Ox and HDC immunohistochemistry staining were performed by automated Ventana BenchMark XT system (Roch, Ventana Medical Systems Inc., Tucson). The first set of sections across the posterior hypothalamus was deparaffinized in xylene and rehydrated through a graded alcohol series. After washing in aquadest (2 x 5 min), slices were briefly washed (3 min) and microwave-treated with 0.01 M sodium citrate in phosphate buffer saline (PBS), pH = 6.0; 2 x 5 minutes) for antigen retrieval. A one-hour pre-incubation step in 5% PB-milk (0.05 M Tris, 0.15 M NaCl, pH=7.6) at room temperature was included to reduce background staining. Sections were incubated in rabbit-IgG, anti- Ox-A (Catalog no. 003-30, Phoenix Pharmaceuticals, Inc., Belmont, CA, USA) at a 1:4000 concentration in supermix-5% milk (0.25% gelatin, 0.5% Triton in PBS) at room temperature (RT) for an hour, followed by overnight incubation at 4°C in the moist chamber. Followed by a one-hour secondary antibody incubation in biotinylated anti-rabbit IgG at a concentration of 1:400 at RT, antigen signal amplification was achieved by a one-hour incubation in ABC-complex (Vectastain®, Avidin-Biotin complex kit). Visualization of the signal was reached by a 20 min incubation of the hypothalamic sections in PBS containing 3.3’-diaminobenzidine (DAB; Sigma) and 0.01% vol/vol H2O2 (Merck, Whitehouse Station, NJ, USA).

The second set of sections underwent exactly the same immunohistochemical procedure using a rabbit-IgG, anti-HDC (EUD 2601, Acris) at a dilution of 1:4000.

Both Ox-A and HDC antibodies stained unambiguously and distinctively posterior hypothalamic neuronal populations. To test the antibody specificity, both Ox-A and HDC antibodies were tested on Ox- and HDC- knockout mouse brains. As already demonstrated in previous studies, the immunolabeling of Ox-A and HDC in the mouse brains was in general dense and clearly contrasted with the background, with well-defined somata, proximal dendrites, and axons approximate to perikaryal. In contrast, incubation with the Ox-A antibody resulted in no staining in the Ox KO brains and that of the HDC one produced no labeling in tuberomammillary nucleus (TMn) and adjacent areas in HDC-KO mice[12], attesting the selectivity of the antibodies used in the human hypothalamic tissue.

##### Cell counts on Ox- and HDC-immunoreactive neurons

Based upon the counterstaining by Nissel, the medial lateral and the TMn borders were roughly defined as the distance between the most rostral and the most caudal section that contained three or more typical TMN neurons as well as the descending position of fornix [19], while the borders were subsequently determined more precisely by immunocytochemistry for Ox-A and HDC. In each section, all immunopositive neurons with their typical cell profiles and a visible nucleolus and developed perikarya were counted using light microscopy at a magnification of 40x. For quantification of cell counts, we subdivided the rostral-caudal length of the TMN regions and the perifornical areas of the dorso- and medial- parts of the lateral hypothalamus into 3 parts, thus distinguishing in 3 sub-regions: rostral, central (where the most numerous labeled cells were found) and caudal parts. These subdivisions were similar to a previously performed arbitrary subdivision for the TMn and hypothalamus[19, 20]. Two experienced researchers did cell counting blind to the nature of the tissue.

##### Statistical analysis CSF study

Fisher’s exact chi-square and Kruskal-Wallis tests were used for descriptive analyses. Mixed models were performed to study the Ox ranks and the histamine ranks in function of age (0-2 months, 2-6 months, 6-12 months) in the three groups of infants (SIDS, ED, Controls). Groups were also compared 2 by 2 and p-values were corrected for multiple comparisons. Statistical analyses were performed using SAS^®^ version 9.4 (SAS Institute Inc, Cary, NC, USA). In all studies, statistical significance was defined with a level of p < 0.05.

##### Immunohistochemical study

The differences between the groups were evaluated by Mann-Whitney U tests. All values are reported as median minimum and maximum values.

### 2.3. Ethics

At the admission, parents were informed of the possible use of the biological samples from their kids for research purposes and they signed a consent form.

## 3. RESULTS

### 3.1. CSF Orexin, histamine and tele-methylhistamine in SIDS, ED and control groups

#### The CSF Orexin levels

No difference in the Ox CSF levels was observed globally between SIDS, ED and controls (p=0.45) or when compared 2 by 2 (all p> 0.47). In addition, no sex difference was found between groups. However, when analyzing by age groups (Table 1 and Figure 1), the 2-6 months group showed higher Ox values for deceased infants (385 pg/ml (range from 179 to 484 pg/ml) for SIDS and 347 pg/ml (range from 234 to 486 pg/ml) for ED) compared to controls 276 pg/ml (from 200 to 673 pg/ml), p=0.003. This was confirmed when tested 2 by 2 (Table 1).

**Figure 1.**
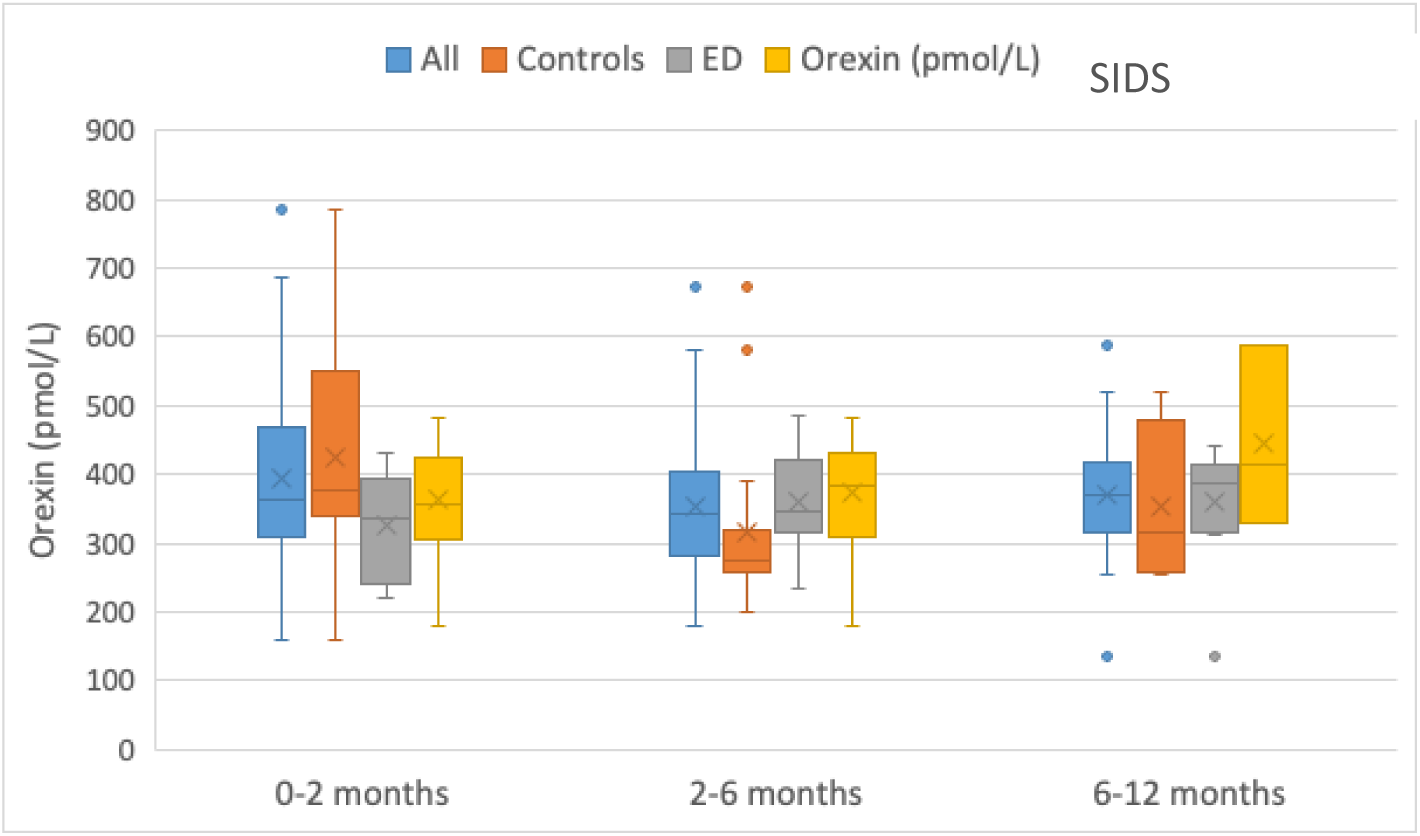
Box plots of the Ox CSF values in all infants, controls, SIDS and infants died of other causes (ED) according to the age (0-2 months, 2-6 months and 6-12 months). The horizontal solid bars represent the Q1-mean-Q3 values, the X is the median, and the vertical line presents the min-max values.

**Table 1.** Demographic characteristics and CSF Ox in healthy controls, SIDS and no-SIDS infants (ED) according to the age (0-2 months, 2-6 months and 6-12 months).

|  | Controls | SIDS | ED | Kruskal Wallis p- values |  |  |  |
| --- | --- | --- | --- | --- | --- | --- | --- |
|  | % (N) or median<br>(min-max) | % (N) or median<br>(min-max) | % (N) or median<br>(min-max) | Global | SIDS vs<br>Controls | ED vs<br>Controls | ED vs<br>SIDS |
| **0-2 months** |  |  |  |  |  |  |  |
| N | 40 | 10 | 9 |  |  |  |  |
| Sex (girls) | 50% (20) † | 50% (5) † | 44% (4) † |  |  |  |  |
| Age (wks) | 4.6 (1.6-10.9) | 5 (2.0-7.0) | 6.7 (1.2-16.4) | 0.41 | 0.79 | 0.44 | 0.66 |
| Corrected age (wks) | 3.6 (0.6-7.9) | 3.9 (0.0-6.0) | 5 (0-7.6) | 0.60 | 0.85 | 0.84 | 0.39 |
| Orexin (pmol/L) | 376 (159-787) | 339.5 (254-466) | 360 (222-430) | 0.12 | 0.98 | 0.29 | 0.23 |
| **≥ 2-6 months** |  |  |  |  |  |  |  |
| N | 17 | 25 | 12 |  |  |  |  |
| Sex (girls) | 47% (8) † | 36% (9) † | 42% (5) † |  |  |  |  |
| Age (wks) | 13.3 (8.4-22.0) | 17.4 (8.0-23.9) | 18.2 (11.1-32.7) | 0.08 | 0.29 | 0.06 | 0.66 |
| Corrected age (wks) | 11.6 (8.4-22.0) | 16.4 (8.0-23.9) | 16.5 (8.1-23.3) | 0.22 | 0.33 | 0.23 | 0.98 |
| Orexin (pmol/L) | 276 (200-673) | 385. (179-484) | 347 (234-486) | 0.003 | 0.003 | 0.04 | 0.79 |
| **≥ 6-12 months** |  |  |  |  |  |  |  |
| N | 4 | 4 | 10 |  |  |  |  |
| Sex (girls) | 50% (2) † | 100% (4) † | 50% (5) † |  |  |  |  |
| Age (wks) | 26.6 (25.0-36.0) | 28.6 (26.0-32.0) | 35.9 (28.7-52.4) | 0.02 | 0.66 | 0.09 | 0.04 |
| Corrected age (wks) | 25.9 (24.4-35) | 25.6 (17.0-32.0) | 33.9 (25.0-48.0) | 0.07 | 0.96 | 0.24 | 0.12 |
| Orexin (pmol/L) | 315.5 (256-519) | 380.5 (328-588) | 386 (135-442) | 0.65 | 0.66 | 0.84 | 0.84 |

When evaluating the maturation of the Ox levels according to age (i.e., within each infant group), a variation of Ox levels with age was only observed in the control group with decreased Ox levels from 0-2 months to 2-6 months of life (p=0.009).

In deceased infants, no difference in CSF Ox levels were observed according to known risk factors for SIDS, such as ventral position, co-sleeping, smoke exposure. However, the CSF Ox levels were higher in died premature (median 407 (225-588) vs 352 (135-473) Kruskal Wallis p=0.03) and in infants died of infection (404 (135- 588) vs 348 (179-484), Kruskal Wallis p=0.003).The duration between infant death and CSF sampling was not different in ED and SIDS infants (median (Q1-Q3) of 3hrs40 (3hrs10-4hrs35) vs 2hrs50 (1hr10-12hrs50), p=0.31). No association was found between delay of lumbar puncture and Ox levels in both groups.

#### The Histamine & tele-methylhistamine levels

Sex differences were observed, globally and when groups were compared 2 by 2 but only in infants aged 2 to 6 months with more girls in the ED group than in the control one and less girls in the SIDS group than in the control one. In addition, age differences were observed within the age categories including infants older than 2 months of age. The same was observed for age in children within age categories over 2 months of age (Table 2, Fig 2A). In mixed models accounting for age and sex, deceased infants always present markedly higher HA levels than controls and SIDS infants also show higher t- MeHA levels (Table 2, Fig 2B) due to a postmortem cytolysis. No association was found between delay of lumbar puncture and HA or t- MeHA levels in ED and SIDS infants.

**Figure 2A.**
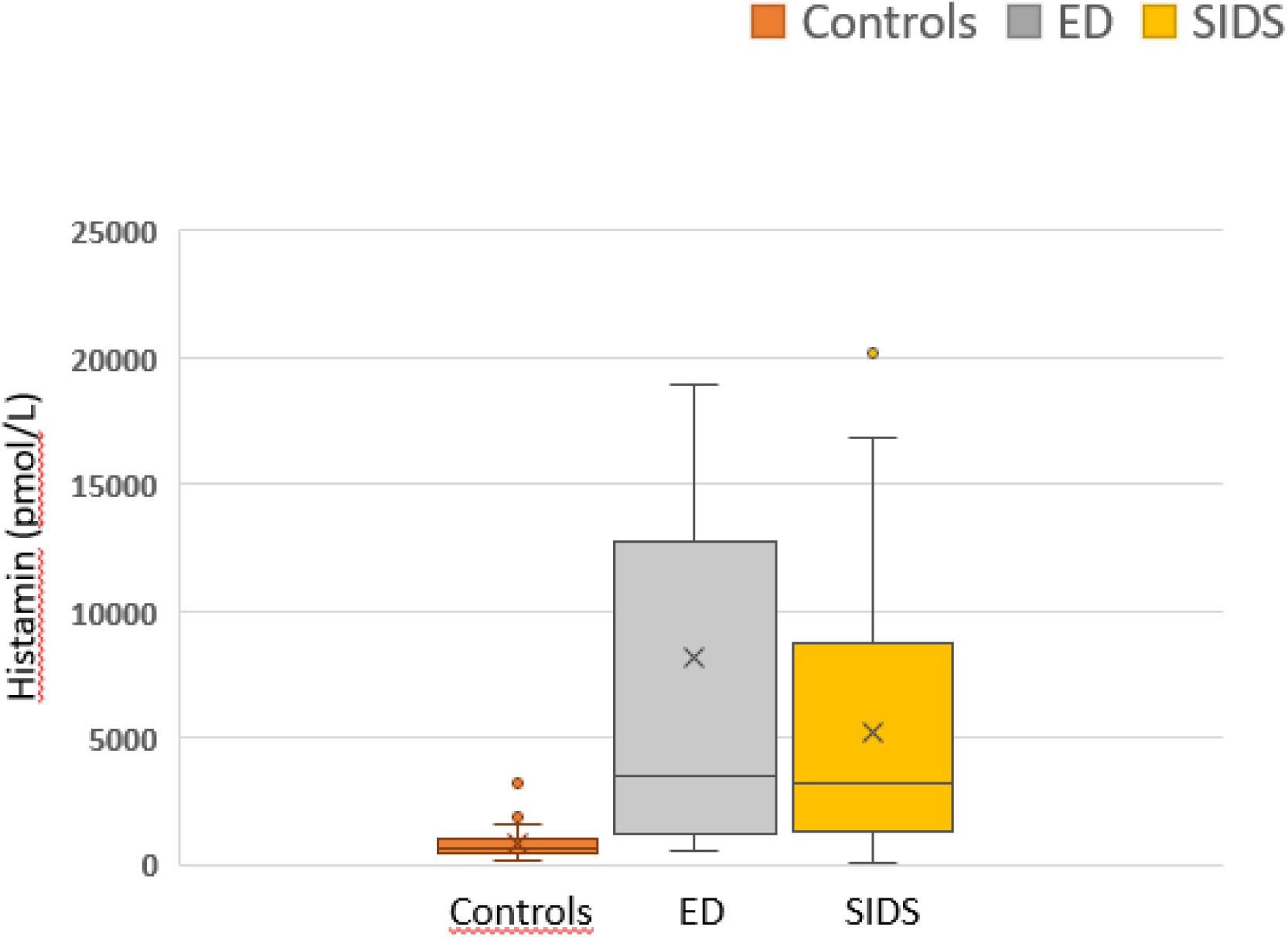
Box plots of CSF Histamine levels in controls, SIDS and infants died of other causes (ED). The horizontal solid bars represent the Q1-mean-Q3 values, the X is the median, and the vertical line presents the min-max values.

**Figure 2B.**
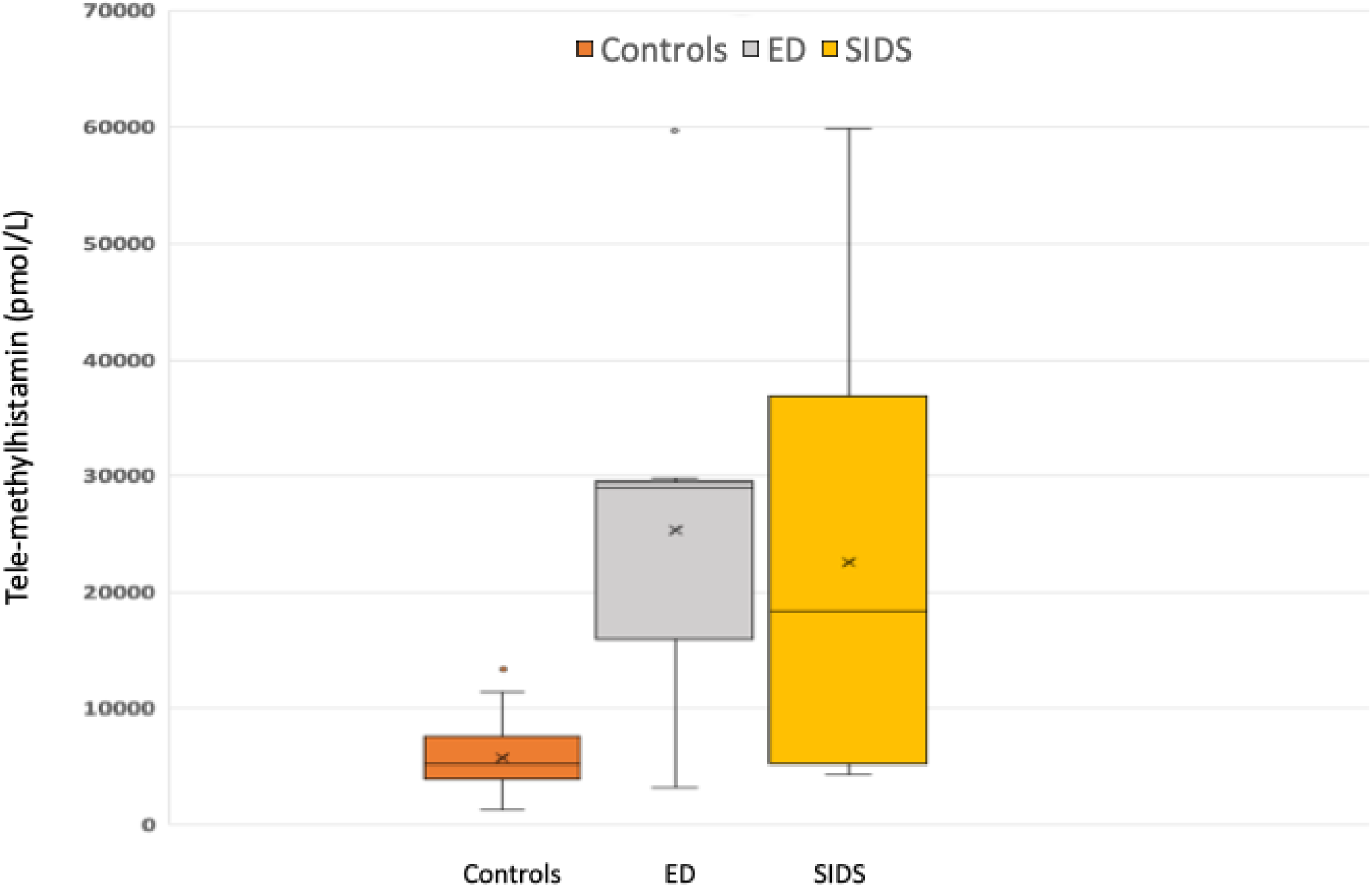
Box plots of CSF Tele-methylhstamine (t-MeHA) levels in controls, SIDS and infants died of other causes (ED). The horizontal solid bars represent the Q1-mean-Q3 values, the X is the median, and the vertical line presents the min-max values.

**Table 2.**
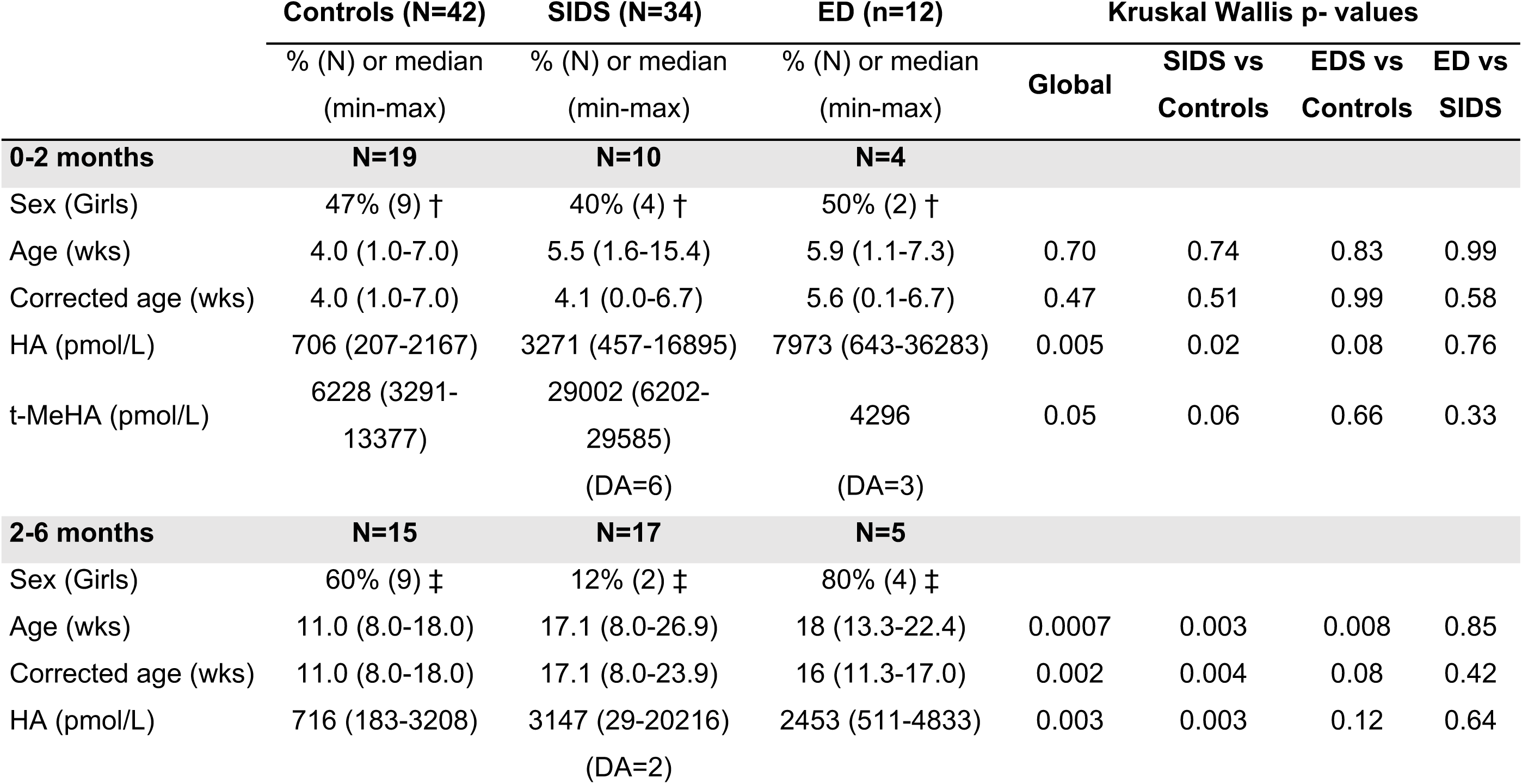
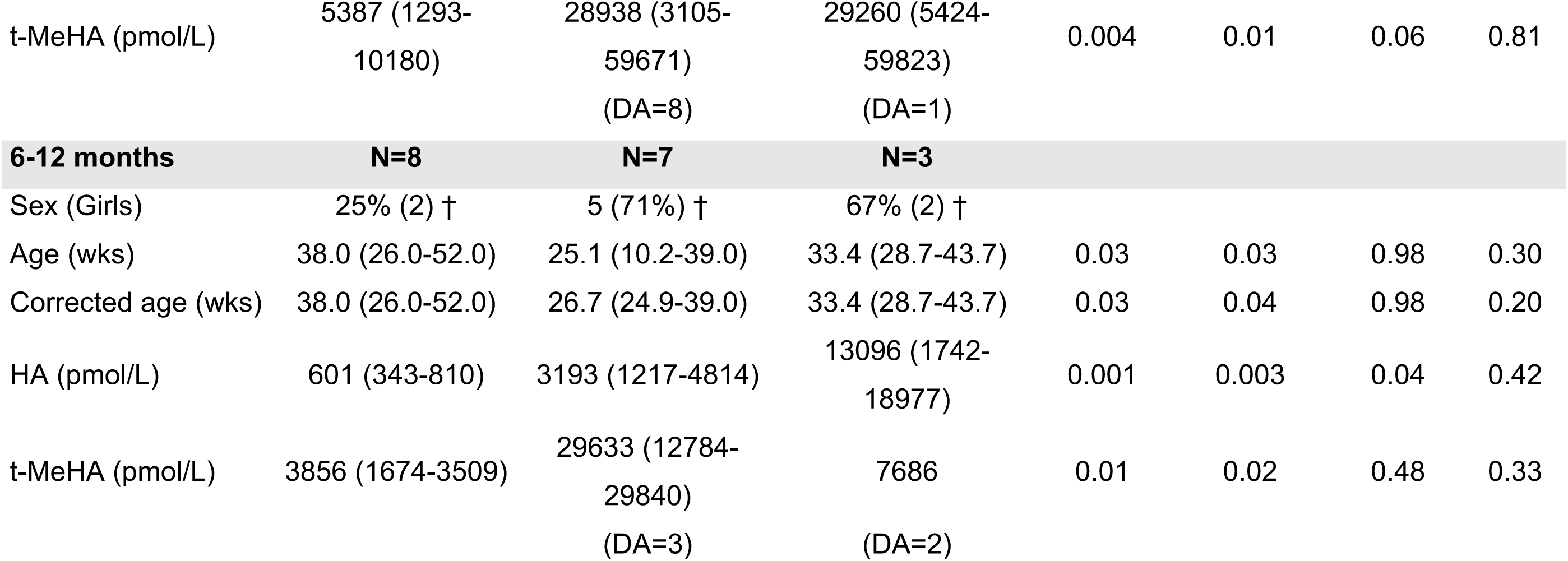
Demographic characteristics and CSF HA and t-MeHA in healthy controls, SIDS and ED infants according to age (0-2 months, 2-6 months and 6-12 months). † Fisher exact test p>0.10 ; ‡ Fisher exact test p=0.003.

### 3.2. Cell counts and morphology of HDC- and Ox-immunoreactive neurons in the postmortem brain tissue

Table 3 shows demographic data for the included infants. There was no difference for age (p Kruskal-Wallis=0.37), sex (p Kruskal-Wallis =1.0), and gestational age (p Kruskal-Wallis =0.14) between SIDS and controls.

**Table 3.**
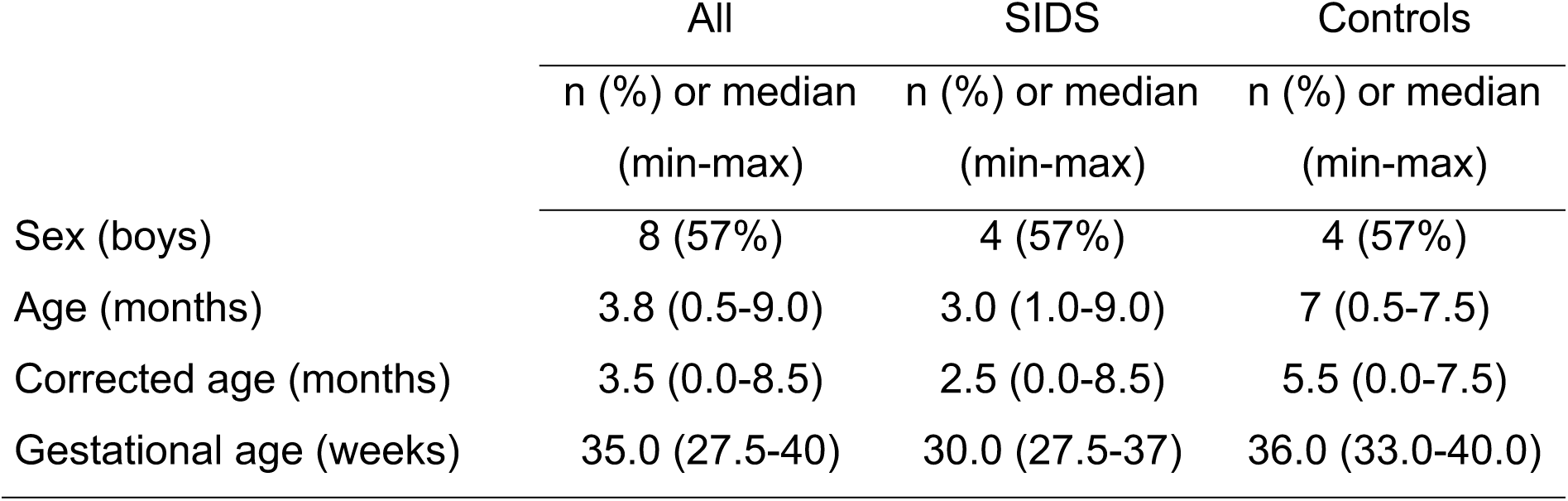
Demographic characteristics of SIDS and no-SIDS infants (ED) for cell counts on Ox- and HA-immunoreactive neurons.

On the control brain sections, numerous HDC-immunoreactive (ir) cell bodies were located in the posterior basal hypothalamus. They were particularly concentrated in the tuberomammillary nucleus (TMn) and adjacent areas which embodied a major part of the ventral posterior hypothalamus. The distribution of HDC-ir perikarya in the SIDS brains was similar to that seen with control ones and no clear distinction could be visualized (Figure 3).

**Figure 3.**
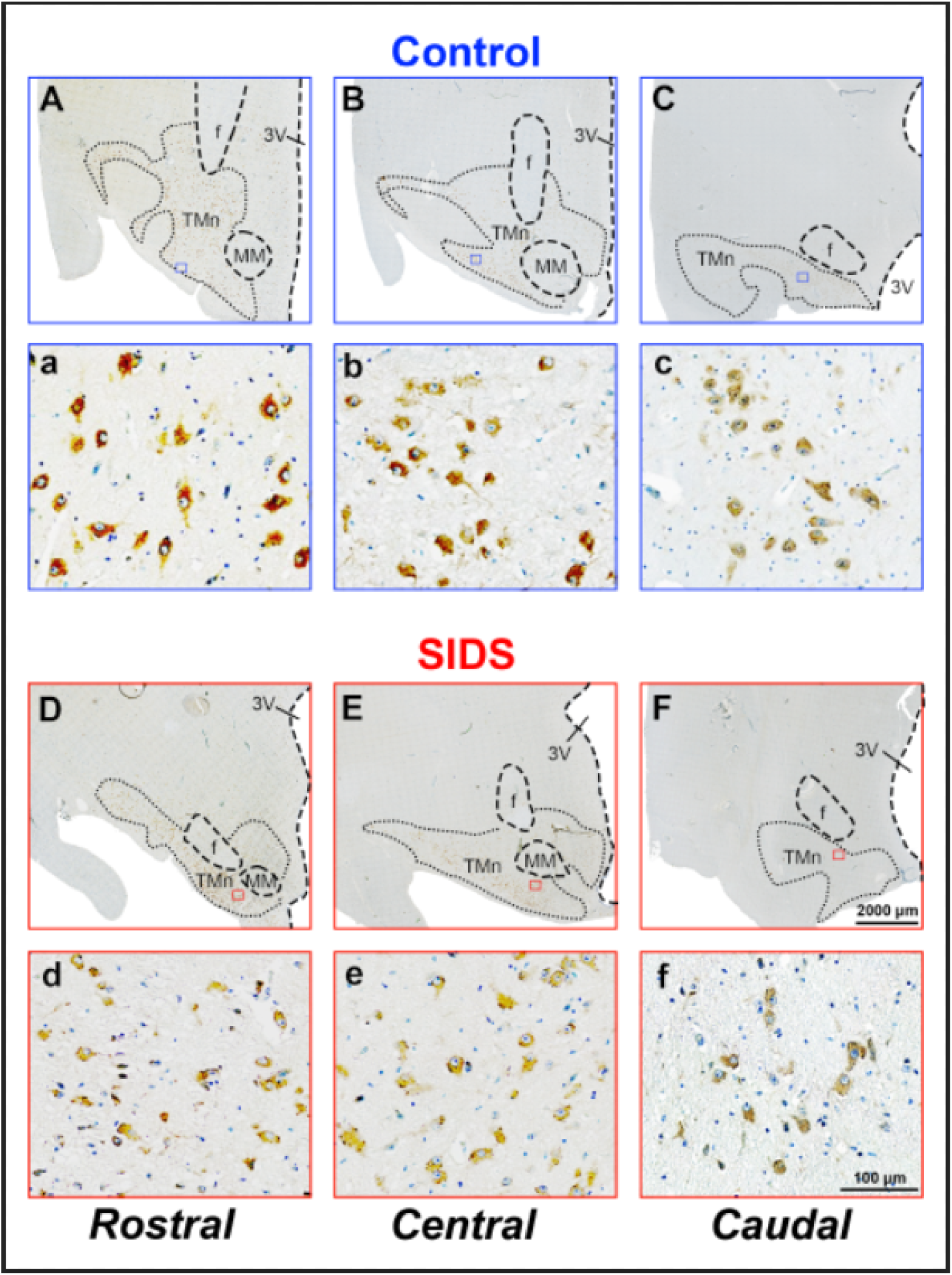
Photomicrographs from representative immunoreacted coronal sections illustrating histidine-decarboxylase (HDC)-immunoreactive neurons across the ventral posterior hypothalamus in control (upper, A-C and a-c) and SIDS (lower, D-F and d-f) groups. Sections were counterstained by Nissel (blue). A-F, lower magnification showing the distribution of stained neurons; a – f, higher magnification from the part of the tuberomammillary nucleus (TMn) indicated by a blue or red inset in A - F, showing the details of the labelling and morphology of stained neurons. Note the similar distribution and morphology of HDC-ir neurons between control and SIDS examples. Abbreviations: DM, dorsal medial hypothalamus; f, fornix; LH, lateral hypothalamus; MM, mammillary nucleus; 3V, third ventricle. Scale bars = 2000 μm in A - F and 100 μm in a - f.

In both control and SIDS brains, Ox-ir cell bodies were located dorsorostral to HDC-ir neurons in the posterior hypothalamus. Their perikarya were concentrated on the upper peri-fornical areas of the dorsal part of the posterior hypothalamus. They were distributed in a more scattered and lesser dense manner than that of HDC-ir neurons. Their distribution and morphology were similar between the control and SIDS brains (Figure 4).

**Figure 4.**
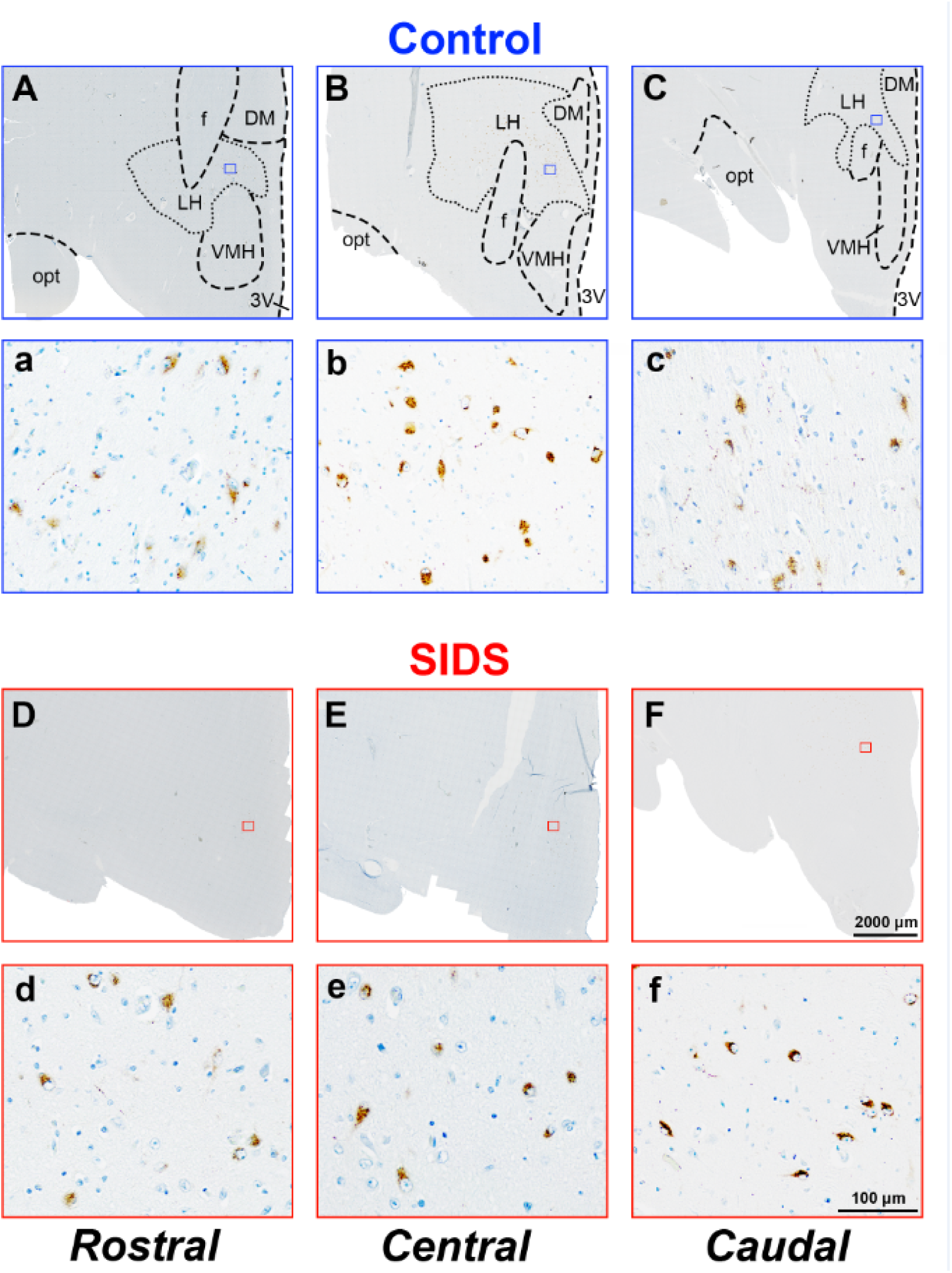
Photomicrographs from representative immunoreacted coronal sections illustrating orexin (Ox) A-immunoreactive neurons across the dorsal part of the posterior hypothalamus in control (upper, A-C and a-c) and SIDS (lower, D-F and d-f) patients. Sections were counterstained by Nissel (blue). A-F, lower magnification showing the distribution of stained neurons; a – f, higher magnification from a blue or red inset in A-F, showing the details of the labelling and morphology of stained neurons in the dorsal part of the posterior hypothalamus. Note the similar distribution and morphology of Ox A-ir neurons between control and SIDS examples. Abbreviations: DM, dorsal medial hypothalamus; f, fornix; LH, lateral hypothalamus; opt, optic tractus; 3V, third ventricle. Scale bars = 2000 μm in A - F and 100 μm in a - f.

When counting the cell number for the multiple brain sections in the same part within some infants, there was no global difference for Ox (P=0.47) or HDC (p=0.14) immunoreactive neurons in SIDS compared to Controls (Table 4).

**Table 4:**
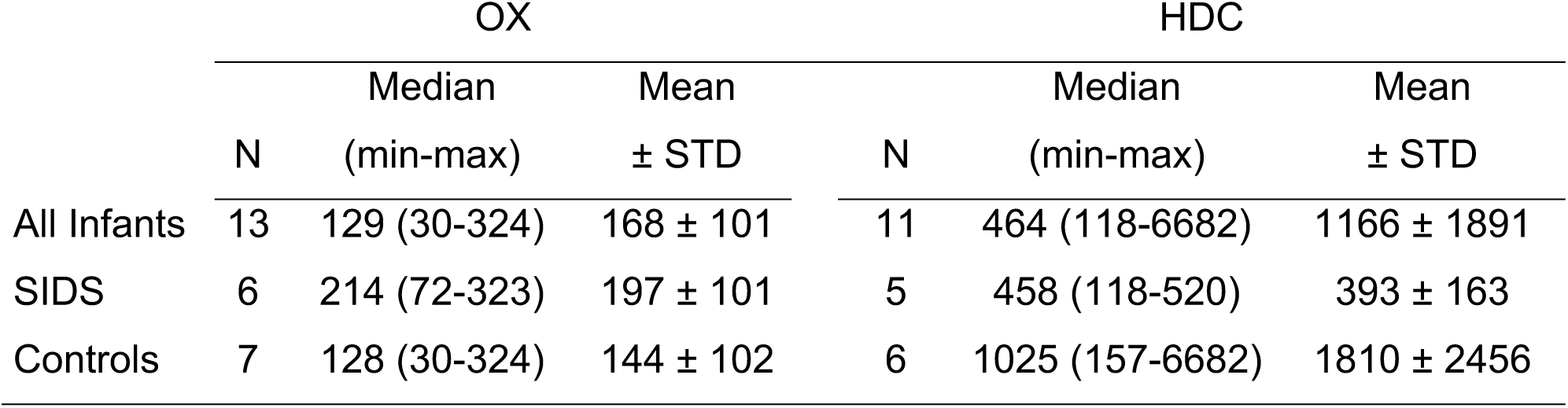
OX and HDC immunoreactive neurons in SIDS and controls.

We then subdivided the HDC- and Ox-ir neuronal populations into rostral, central and caudal parts in order to quantify their respective cell numbers and then compare them between SIDS and control subjects.

Regarding HDC-ir neurons, a high variability was observed in their cell counts in Controls compared to SIDS infants, especially in the central and rostral parts (Figure 5). No significant difference was found between SIDS and controls in rostral (p=1.0), central (p=0.67) and caudal (p=0.45) parts. In addition, microscopic observation did not reveal any detectable difference between the two groups in terms of neuronal morphology (Fig 3).

**Figure 5:**
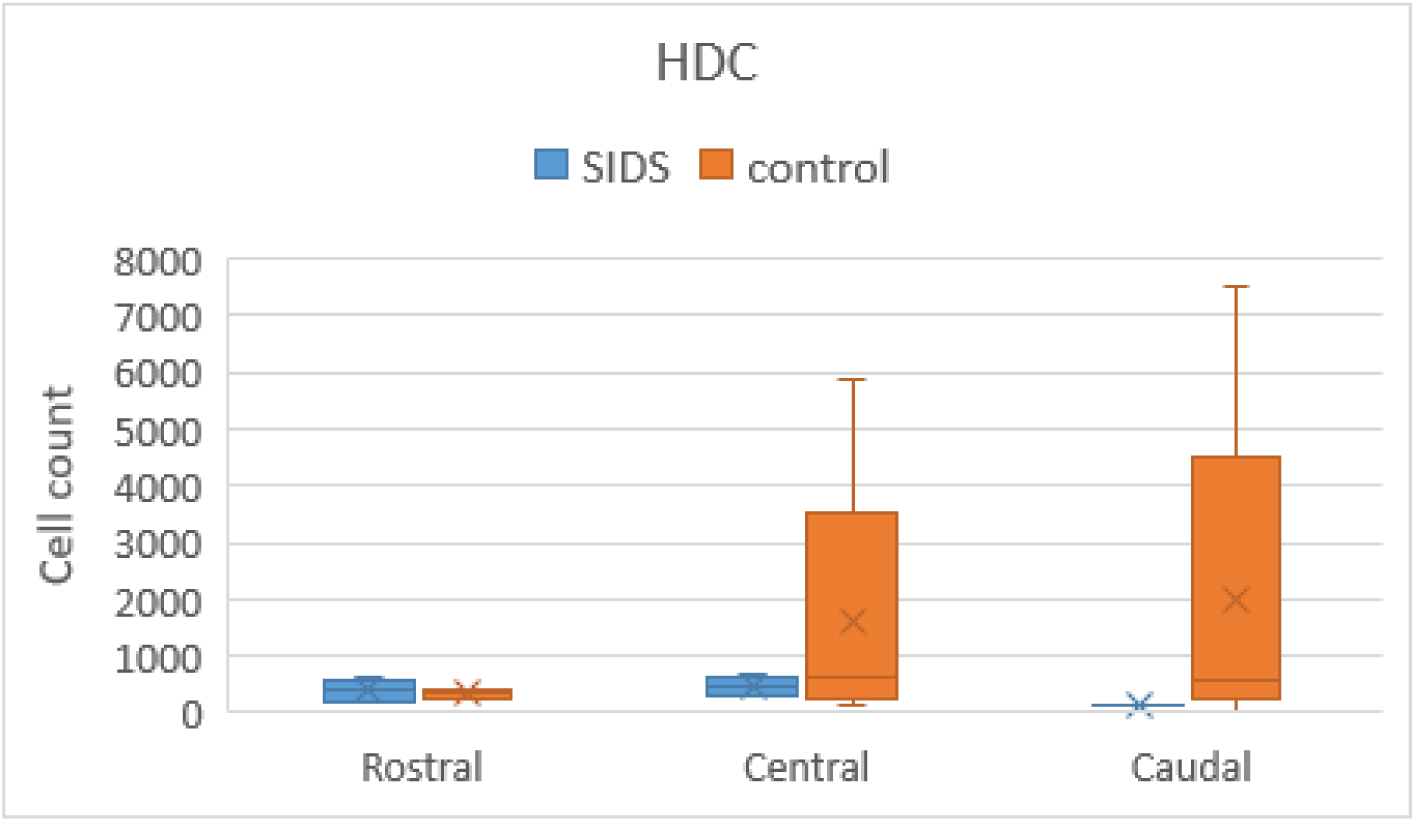
Box plots showing cell counts and quantification of the number of histidine-decarboxylase (HDC)- immunoreactive (ir) neurons from the rostral, central and caudal subdivisions of the tuberomammillary nucleus and adjacent areas in control and SIDS subjects. The horizontal solid bars represent the Q1-mean-Q3 values, the X is the median, and the vertical line presents the min-max values.

Regarding Ox-ir neurons, SIDS infants show higher cell number in rostral (Kruskall Wallis p=0.04) and caudal (p=0.03) subdivisions than in controls but not in central subdivision (p=0.60) (Figure 6). No difference was observed in terms of their morphology (Figure 4).

**Figure 6.**
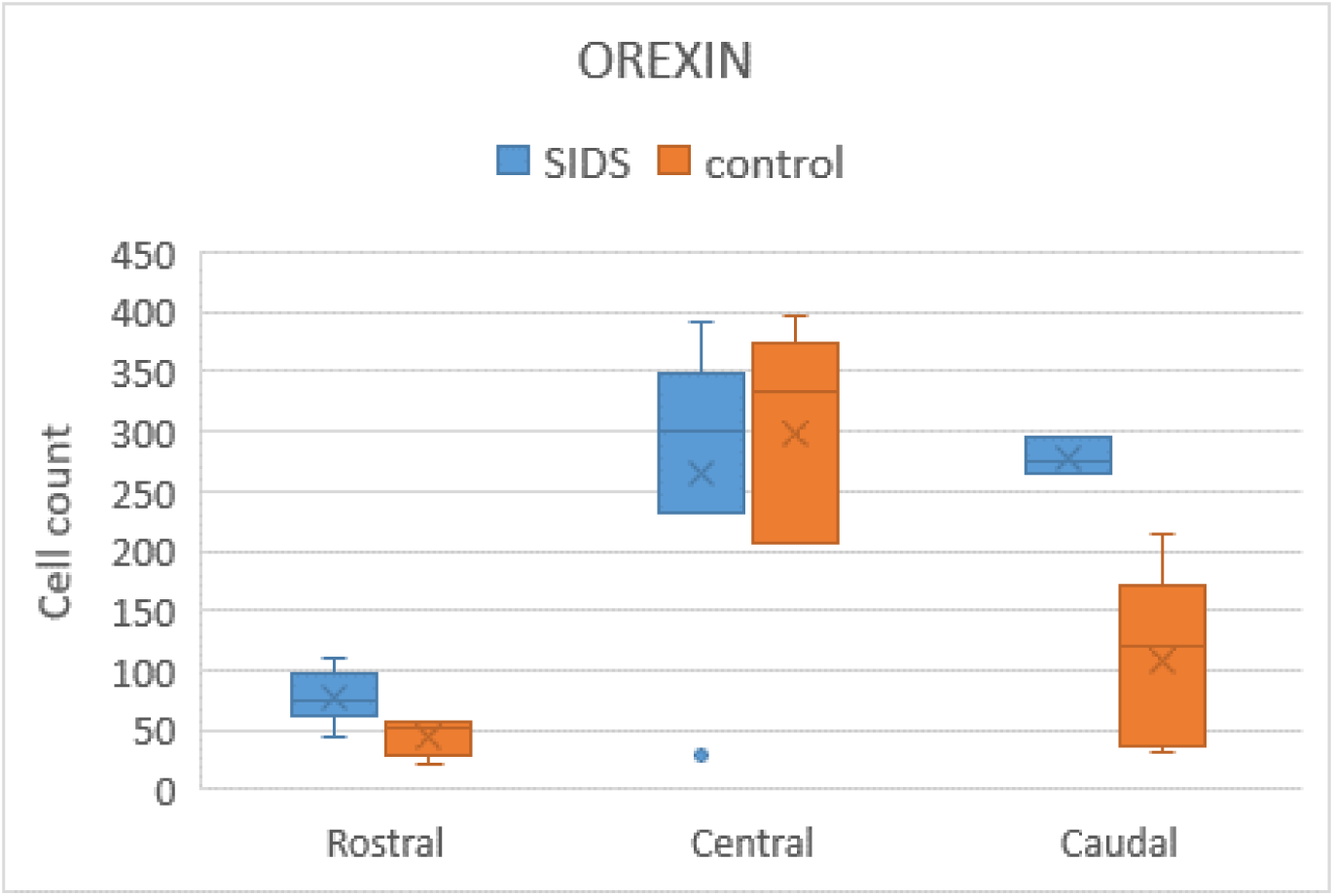
Box plots showing cell counts and quantification of the number of orexin A-immunoreactive (ir) neurons from the rostral, central and caudal subdivisions of the lateral hypothalamus in control and SIDS subjects. The horizontal solid bars represent the Q1-mean-Q3 values, the X is the median, and the vertical line presents the min-max values.

## 4. DISCUSSION

Sudden infant death syndrome (SIDS), characterized by dysfunctional sleep and impaired arousal thresholds, is generally believed to be associated with disrupted arousal mechanisms and vulnerable respiratory control systems in the brain [21, 22]. Supporting this hypothesis, neurochemical abnormalities have been reported in brain regions involved in sleep regulation, arousal, and responses to hypoxia and/or hypercapnia—conditions frequently linked to prone sleeping and viral infections, both known as SIDS risk factors [1, 23–26].

Early investigations into the pathogenesis of SIDS in both humans and animal models have primarily focused on the brainstem, particularly the serotonergic raphe system [27]. Since the early 2000s, the serotonergic system has been implicated through findings of decreased serotonin metabolite (5-HIAA) levels in the serotonergic nuclei of the medulla (including the median raphe) in approximately 30% of SIDS cases[24]. More recently Harrington et al. identified a possible involvement for the cholinergic system [28]. SIDS infants showed significantly lower butyrylcholinesterase (BChE) activity compared to both non-SIDS deceased infants and age-matched living controls. These results suggest autonomic nervous system dysfunction, particularly parasympathetic dysregulation, which has been supported by electrophysiological studies [29–31].

Given these findings, it is essential to further explore other brain structures, neurotransmitters, and neuropeptides involved in sleep and arousal. In this context, we turned our attention to the hypothalamic wake-promoting systems-specifically histamine and orexin (hypocretin) neurons—which are known to contribute to cortical and behavioral arousal during wakefulness [11, 12, 32]. These raise critical questions: Are orexin and histamine transmissions implicated in SIDS pathogenesis? Are they involved via similar or independent mechanisms? Could they represent hypothalamic biomarkers for SIDS?

We then measured CSF orexin levels in SUDI (SIDS and non-SIDS deaths) and compared them with healthy controls, while analyzing correlations with known SIDS risk factors (e.g., prone sleeping, smoke exposure in utero, infection, prematurity). CSF orexin levels were elevated in both SIDS and non-SIDS infants compared to controls during the 2–6 months age window, consistent with the age of greatest SIDS vulnerability, with particularly high levels in preterm infants and those suffering from infections. Importantly, when comparing overall CSF orexin levels across all SIDS, non-SIDS, and control groups, no statistically significant differences were observed. However, as orexin is a neuropeptide, its CSF concentration is considered relatively stable and not significantly altered postmortem [33]. Moreover, no correlation was found between the postmortem interval and CSF orexin levels.

In addition, we compared the number of orexin-A immunoreactive neurons in the hypothalamus between SIDS and non-SIDS brains. Our findings showed that SIDS victims had a significantly higher number of orexin-A immunoreactive neurons in both the rostral and caudal subdivisions, but not in the central region, compared to infants who died from other causes.

We hypothesize that during the critical 2–6 month age window—when CSF orexin levels are generally lower—infants who succumbed to SUDI may have faced acute stress or hypoxic events (in the case of ED), whereas SIDS victims might have also experienced chronic or prolonged challenges. This could help explain why preterm infants and those exposed to infections, both of whom may have a heightened stress response, displayed elevated CSF orexin levels.

Indeed, an increased activity in some orexin neurons may result from repeated hypoxic episodes prior to death, as suggested by animal studies [34]. The elevated number of orexin-expressing neurons observed in infants who died from SIDS, compared to non- SIDS cases, may reflect chronic hypoxia. This hypoxia could stem from newborns’ postural upper airway obstruction or exposure to other established risk factors such as infection or prematurity, both of which heighten the risk of hypoxic and hypercapnic episodes [1]. Indeed, infants who later succumbed to SIDS were often found to experience more frequent obstructive sleep apneas than healthy controls [5, 35]. Obstructive breathing events may arise from various mechanisms, including anatomical features of the face or airways [36, 37], or abnormalities in the neuronal control of airway patency [38]. Several known SIDS risk factors—such as prematurity, prenatal exposure to tobacco smoke, and upper airway viral infections—have also been associated with an increased incidence of obstructive apneas in infants [39–41].

By their projections to of the solitary tract, hypoglossal nucleus, Kölliker–Fuse nucleus, and phrenic motor nucleus, orexin neurons are well-positioned to regulate respiratory control [42–44]. In vitro data suggest orexin participates in respiratory control during early postnatal life. For instance, OX1R blockade reduces hypoglossal burst frequency in neonatal mouse brainstem slices [45], while exogenous orexin increases cervical spinal nerve activity [46]. In vivo, orexin maintains eupneic respiratory frequency in two-week-old rats, possibly through modulation of peripheral chemoreflex activity [47]. Using plethysmography and EEG/EMG monitoring, Spinelli et al. showed that OXR antagonism reduced ventilatory and blood pressure responses to hypercapnia during the inactive phase of circadian cycle, when hypoxia also activates a subset of perifornical orexin neurons [44]. These results indicate orexin’s role in modulating both central and peripheral chemoreflexes in a state-dependent manner.

However, the literature is not entirely consistent. Hypoxia and hypercapnia can suppress orexin expression rapidly. For instance, adult rats exposed to intermittent hypoxia (6–8% O₂) or hypercapnia (10% CO₂) show a 50–60% decrease in prepro- orexin mRNA levels [48], and neonatal piglets exposed to hypercapnic hypoxia exhibit a 25% reduction in hypothalamic orexin immunoreactivity, without neuronal loss [49]. Animal studies also highlight the complexity of orexin dynamics: intermittent hypoxia produces more pronounced effects than sustained hypoxia [34], responses vary with age [47], circadian phase [44], and regional nuclei [44], and interactions with other arousal systems further modulate these effects [10].

Similarly, Hunt et al. found reduced orexin immunoreactivity in the hypothalamus (−21%) and pons (−40–60%) of SIDS victims, without decrease in cell number [10]. The SIDS victims in that study were notably older (mean age ∼14 months), with only 16% born preterm compared to 36% in controls. These findings may reflect age-related patterns. Previous rodent studies have shown a modest (10–25%) reduction in orexin neuron number, but a much larger (60–80%) drop in fiber immunoreactivity, suggesting increased axonal release prior to death [50]. This phenomenon may apply to SIDS cases as well, where prolonged stress or hypoxia could trigger orexin release and subsequent downregulation.

Given these complexities, we also examined histaminergic neuronal number and CSF levels. Although histaminergic neurons are highly sensitive to hypoxia [51], hypercapnia [52], and arousal failure [53], we found no significant differences in the number of HDC-immunoreactive neurons between SIDS and control cases across hypothalamic subregions. Regarding CSF levels of histamine and its metabolite, both show circadian variation, fluctuate across the sleep–wake cycle under normal conditions. They are altered in conditions such as childhood narcolepsy or excessive sleepiness, supporting a predominantly neuronal origin [14, 54–56]. Under healthy conditions, the contribution of mast cells appears minimal. However, in pathological contexts such as inflammation, infection, or allergic responses, mast cell degranulation can markedly elevate CSF histamine levels [57, 58].

Finally, unlike the neuropeptide orexins, which remain relatively stable after death, postmortem levels of amines such as histamine are subject to alteration. Although we observed significantly higher CSF levels of histamine and tele-methylhistamine in SIDS infants compared to healthy controls, it remains unclear whether these elevations result from increased neuronal release or mast cell degranulation[59–61], and whether they contribute to the pathogenesis of SIDS or merely reflect postmortem changes. Nevertheless, the markedly elevated levels of tele-methylhistamine argue in favor of a neuronal origin, as neuronal histamine is characterized by rapid metabolism and high turnover, driven by highly dynamic HDC activity—unlike mast cells, which exhibit minimal HDC activity and low metabolic rates[62].

## Limitations and Future Directions

Several methodological limitations should be acknowledged. First, the number of postmortem samples was relatively small. Nevertheless, this is the first study to report both orexin/histamine cell counts and CSF levels in the same context, though not always in the same individuals. Ideally, paired data would strengthen correlations. Second, while the HDC antibody used is highly specific in the rat, cross-reactivity with other monoaminergic neurons cannot be excluded in other species [63]. Importantly, this did not confound cell counts in the posterior hypothalamus [64, 65], and the antibody does not stain tuberomammillary neurons in HDC knockout mice. The morphology and distribution of histaminergic neurons in SIDS were comparable to previous studies using HA-specific antibodies in human brains [66]. Future studies with optimized histological fixation and specific HA markers are warranted. Finally, orexin and histamine were not measured in the same individuals, Indeed, due to the limited volume of CSF obtained, it was not possible to measure both of them from the same tube. For practical reasons, it was impossible to obtain both CSF samples and postmortem brain tissue from the same subject. Moreover, larger sample sizes will be necessary. The integration of neurochemical, genetic, and imaging data (e.g., via the Biominrisk project, NCT06244433) [67] will also be essential to understanding the pathophysiology of SIDS.

## 5. CONCLUSIONS

Despite a dramatic decline in SIDS incidence following public health campaigns, it remains the leading cause of post-neonatal mortality. No reliable biomarker currently exists to predict SIDS risk. Although no dysfunction of these systems was directly demonstrated, the elevated orexin cell counts and CSF levels in SIDS infants suggest increased orexin release, potentially due to prolonged or repeated stressors such as hypoxia. Current data on histamine neurons do not support their direct involvement in SIDS, highlighting the need for more sensitive biomarkers.

## Acknowledgements

This study was supported by Laboratory of Integrative Physiology of the Brain Arousal System, INSERM U628 and U1028, Université Claude Bernard Lyon1, ANR NarConX, ANR Histawake and the Talent-Introducing Project of Foreign Experts Affairs of China (X2017008). The work involved GF Cui and YP Hou was supported by National Natural Science Foundation of China (81771426). We thank Pr. Dick F. Swaab for advice, transmitted by Dr. Ling Shan who also helped us in the postmortem histological study of this work by technical and scientific discussion.

## Reference

1. Moon, R.Y., R.S. Horne, and F.R. Hauck, Sudden infant death syndrome. Lancet, 2007. 370(9598): p. 1578–87.

2. de Visme, S., et al., National Variations in Recent Trends of Sudden Unexpected Infant Death Rate in Western Europe. J Pediatr, 2020. 226: p. 179–185 e4.

3. Blair, P.S., et al., Major epidemiological changes in sudden infant death syndrome: a 20-year population-based study in the UK. Lancet, 2006. 367(9507): p. 314–9.

4. Schechtman, V.L., et al., Sleep state organization in normal infants and victims of the sudden infant death syndrome. Pediatrics, 1992. 89(5 Pt 1): p. 865–70.

5. Kahn, A., et al., Sleep and cardiorespiratory characteristics of infant victims of sudden death: a prospective case-control study. Sleep, 1992. 15(4): p. 287–92.

6. Kato, I., et al., Incomplete arousal processes in infants who were victims of sudden death. Am J Respir Crit Care Med, 2003. 168(11): p. 1298–303.

7. Mieda, M. and T. Sakurai, Overview of orexin/hypocretin system. Prog Brain Res, 2012. 198: p. 5–14.

8. Lancien, M., et al., Low cerebrospinal fluid hypocretin levels during sudden infant death syndrome (SIDS) risk period. Sleep Med, 2017. 33: p. 57–60.

9. Aran, A., et al., CSF levels of hypocretin-1 (orexin-A) peak during early infancy in humans. Sleep, 2012. 35(2): p. 187–91.

10. Hunt, N.J., et al., Decreased orexin (hypocretin) immunoreactivity in the hypothalamus and pontine nuclei in sudden infant death syndrome. Acta Neuropathol, 2015. 130(2): p. 185–98.

11. Haas, H.L., O.A. Sergeeva, and O. Selbach, Histamine in the nervous system. Physiol Rev, 2008. 88(3): p. 1183–241.

12. Anaclet, C., et al., Orexin/hypocretin and histamine: distinct roles in the control of wakefulness demonstrated using knock-out mouse models. J Neurosci, 2009. 29(46): p. 14423–38.

13. Parmentier, R., et al., Anatomical, physiological, and pharmacological characteristics of histidine decarboxylase knock-out mice: evidence for the role of brain histamine in behavioral and sleep-wake control. J Neurosci, 2002. 22(17): p. 7695–711.

14. Franco, P., et al., Impaired histaminergic neurotransmission in children with narcolepsy type 1. CNS Neurosci Ther, 2019. 25(3): p. 386–395.

15. Dauvilliers, Y., et al., Normal cerebrospinal fluid histamine and tele-methylhistamine levels in hypersomnia conditions. Sleep, 2012. 35(10): p. 1359–66.

16. Plancoulaine, S., et al., Cerebrospinal Fluid Histamine Levels in Healthy Children and Potential Implication for SIDS: Observational Study in a French Tertiary Care Hospital. Front Pediatr, 2022. 10: p. 819496.

17. Barateau, L., et al., Cerebrospinal fluid monoamine levels in central disorders of hypersomnolence. Sleep, 2021. 44(7).

18. Croyal, M., et al., Histamine and tele-methylhistamine quantification in cerebrospinal fluid from narcoleptic subjects by liquid chromatography tandem mass spectrometry with precolumn derivatization. Anal Biochem, 2011. 409(1): p. 28–36.

19. Shan, L., A.M. Bao, and D.F. Swaab, The human histaminergic system in neuropsychiatric disorders. Trends Neurosci, 2015. 38(3): p. 167–77.

20. Nakamura, S., et al., Loss of large neurons and occurrence of neurofibrillary tangles in the tuberomammillary nucleus of patients with Alzheimer’s disease. Neurosci Lett, 1993. 151(2): p. 196–9.

21. Franco, P., et al., Arousal from sleep mechanisms in infants. Sleep Med. 11(7): p. 603–14.

22. Thach, B.T., The role of respiratory control disorders in SIDS. Respir Physiol Neurobiol, 2005. 149(1-3): p. 343–53.

23. Kinney, H.C., et al., Subtle autonomic and respiratory dysfunction in sudden infant death syndrome associated with serotonergic brainstem abnormalities: a case report. J Neuropathol Exp Neurol, 2005. 64(8): p. 689–94.

24. Duncan, J.R., et al., Brainstem serotonergic deficiency in sudden infant death syndrome. JAMA, 2010. 303(5): p. 430–7.

25. Randall, B.B., et al., Potential asphyxia and brainstem abnormalities in sudden and unexpected death in infants. Pediatrics, 2013. 132(6): p. e1616–25.

26. Waters, K.A., N.J. Hunt, and R. Machaalani, Neuropathology of Sudden Infant Death Syndrome: Hypothalamus, in SIDS Sudden Infant and Early Childhood Death: The Past, the Present and the Future, J.R. Duncan and R.W. Byard, Editors. 2018: Adelaide (AU).

27. Kinney, H.C. and B.T. Thach, The sudden infant death syndrome. N Engl J Med, 2009. 361(8): p. 795–805.

28. Harrington, C.T., N.A. Hafid, and K.A. Waters, Butyrylcholinesterase is a potential biomarker for Sudden Infant Death Syndrome. EBioMedicine, 2022. 80: p. 104041.

29. Kluge, K.A., et al., Spectral analysis assessment of respiratory sinus arrhythmia in normal infants and infants who subsequently died of sudden infant death syndrome. Pediatr Res, 1988. 24(6): p. 677–82.

30. Franco, P., et al., Decreased autonomic responses to obstructive sleep events in future victims of sudden infant death syndrome. Pediatr Res, 1999. 46(1): p. 33–9.

31. Franco, P., et al., Polysomnographic study of the autonomic nervous system in potential victims of sudden infant death syndrome. Clin Auton Res, 1998. 8(5): p. 243–9.

32. Lin, J.S., O.A. Sergeeva, and H.L. Haas, Histamine H3 receptors and sleep-wake regulation. J Pharmacol Exp Ther, 2011. 336(1): p. 17–23.

33. Peyron, C., et al., A mutation in a case of early onset narcolepsy and a generalized absence of hypocretin peptides in human narcoleptic brains. Nat Med, 2000. 6(9): p. 991–7.

34. Yamaguchi, K., et al., Intermittent but not sustained hypoxia activates orexin-containing neurons in mice. Respir Physiol Neurobiol, 2015. 206: p. 11–4.

35. Kato, I., et al., Developmental characteristics of apnea in infants who succumb to sudden infant death syndrome. Am J Respir Crit Care Med, 2001. 164(8 Pt 1): p. 1464–9.

36. Ducloyer, M., et al., The Ogival Palate: A New Risk Marker of Sudden Unexpected Death in Infancy? Front Pediatr, 2022. 10: p. 809725.

37. Guilleminault, C., et al., Obstructive sleep apnea and near miss for SIDS: I. Report of an infant with sudden death. Pediatrics, 1979. 63(6): p. 837–43.

38. Weese-Mayer, D.E., et al., Association of the serotonin transporter gene with sudden infant death syndrome: a haplotype analysis. Am J Med Genet A, 2003. 122(3): p. 238–45.

39. Kahn, A., et al., Prenatal exposure to cigarettes in infants with obstructive sleep apneas. Pediatrics, 1994. 93(5): p. 778–83.

40. Blackwell, C., Infection, inflammation and SIDS. FEMS Immunol Med Microbiol, 2004. 42(1): p. 1–2.

41. Huang, Y.S., et al., Sleep-disordered breathing, craniofacial development, and neurodevelopment in premature infants: a 2-year follow-up study. Sleep Med, 2019. 60: p. 20–25.

42. Fung, S.J., et al., Hypocretin (orexin) input to trigeminal and hypoglossal motoneurons in the cat: a double-labeling immunohistochemical study. Brain Res, 2001. 903(1-2): p. 257–62.

43. Yokota, S., et al., Orexinergic fibers are in contact with Kolliker-Fuse nucleus neurons projecting to the respiration-related nuclei in the medulla oblongata and spinal cord of the rat. Brain Res, 2016. 1648(Pt A): p. 512–523.

44. Spinieli, R.L., et al., Orexin facilitates the ventilatory and behavioral responses of rats to hypoxia. Am J Physiol Regul Integr Comp Physiol, 2022. 322(6): p. R581–R596.

45. Corcoran, A.E., G.B. Richerson, and M.B. Harris, Functional link between the hypocretin and serotonin systems in the neural control of breathing and central chemosensitivity. J Neurophysiol, 2015. 114(1): p. 381–9.

46. Sugita, T., et al., Orexin induces excitation of respiratory neuronal network in isolated brainstem spinal cord of neonatal rat. Respir Physiol Neurobiol, 2014. 200: p. 105–9.

47. Spinieli, R.L., et al., Orexin contributes to eupnea within a critical period of postnatal development. Am J Physiol Regul Integr Comp Physiol, 2021. 321(4): p. R558–R571.

48. Qiao, Y., et al., Effect of intermittent hypoxia on neuro-functional recovery post brain ischemia in mice. J Mol Neurosci, 2015. 55(4): p. 923–30.

49. Du, M.K., et al., Cumulative effects of repetitive intermittent hypercapnic hypoxia on orexin in the developing piglet hypothalamus. Int J Dev Neurosci, 2016. 48: p. 1–8.

50. Hunt, N.J., et al., Changes in orexin (hypocretin) neuronal expression with normal aging in the human hypothalamus. Neurobiol Aging, 2015. 36(1): p. 292–300.

51. Fan, Y.Y., et al., Activation of the central histaminergic system is involved in hypoxia-induced stroke tolerance in adult mice. J Cereb Blood Flow Metab, 2011. 31(1): p. 305–14.

52. Kernder, A., et al., Acid-sensing hypothalamic neurons controlling arousal. Cell Mol Neurobiol, 2014. 34(6): p. 777–89.

53. Lozeva, V., et al., Increased concentrations of histamine and its metabolite, tele-methylhistamine and down-regulation of histamine H3 receptor sites in autopsied brain tissue from cirrhotic patients who died in hepatic coma. J Hepatol, 2003. 39(4): p. 522–7.

54. Mochizuki, M., et al., Effects of smoking on fetoplacental-maternal system during pregnancy. Am J Obstet Gynecol, 1984. 149(4): p. 413–20.

55. Zeitzer, J.M., et al., Time-course of cerebrospinal fluid histamine in the wake-consolidated squirrel monkey. J Sleep Res, 2012. 21(2): p. 189–94.

56. Kanbayashi, T., et al., CSF histamine contents in narcolepsy, idiopathic hypersomnia and obstructive sleep apnea syndrome. Sleep, 2009. 32(2): p. 181–7.

57. Rozniecki, J.J., et al., Elevated mast cell tryptase in cerebrospinal fluid of multiple sclerosis patients. Ann Neurol, 1995. 37(1): p. 63–6.

58. Letourneau, R., et al., Ultrastructural evidence of brain mast cell activation without degranulation in monkey experimental allergic encephalomyelitis. J Neuroimmunol, 2003. 145(1-2): p. 18–26.

59. Edston, E., et al., Increased mast cell tryptase in sudden infant death - anaphylaxis, hypoxia or artefact? Clin Exp Allergy, 1999. 29(12): p. 1648–54.

60. Holgate, S.T., et al., The anaphylaxis hypothesis of sudden infant death syndrome (SIDS): mast cell degranulation in cot death revealed by elevated concentrations of tryptase in serum. Clin Exp Allergy, 1994. 24(12): p. 1115–22.

61. Bonelli, A., et al., Immunohistochemical localization of mast cells as a tool for the discrimination of vital and postmortem lesions. Int J Legal Med, 2003. 117(1): p. 14–8.

62. Schwartz, J.C., et al., Histaminergic transmission in the mammalian brain. Physiol Rev, 1991. 71(1): p. 1–51.

63. Mizuguchi, H., et al., Immuno-cross-reactivity of histidine and dopa decarboxylases. Biochem Biophys Res Commun, 1990. 173(3): p. 1299–303.

64. John, J., et al., Greatly increased numbers of histamine cells in human narcolepsy with cataplexy. Ann Neurol, 2013. 74(6): p. 786–93.

65. Valko, P.O., et al., Increase of histaminergic tuberomammillary neurons in narcolepsy. Ann Neurol, 2013. 74(6): p. 794–804.

66. Panula, P., et al., A histamine-containing neuronal system in human brain. Neuroscience, 1990. 34(1): p. 127–32.

67. Ducloyer, M., et al., Identification of novel genetic, neurobiological and radio-anatomical biomarkers for risk stratification of sudden unexpected death in infancy and early childhood: the BIOMINRISK study protocol. BMJ Open, 2025. 15(7): p. e101811.

